# Energy allocation theory for bacterial growth control in and out of steady state

**DOI:** 10.1101/2024.01.09.574890

**Authors:** Arianna Cylke, Diana Serbanescu, Shiladitya Banerjee

**Affiliations:** Department of Physics, Carnegie Mellon University, Pittsburgh, PA 15213, USA; Department of Physics and Astronomy, University College London, London WC1E 6BT, UK; Institute for the Physics of Living Systems, University College London, London WC1E 6BT, UK

## Abstract

Efficient allocation of energy resources to key physiological functions allows living organisms to grow and thrive in diverse environments and adapt to a wide range of perturbations. To quantitatively understand how unicellular organisms utilize their energy resources in response to changes in growth environment, we introduce a theory of dynamic energy allocation which describes cellular growth dynamics based on partitioning of metabolizable energy into key physiological functions: growth, division, cell shape regulation, energy storage and loss through dissipation. By optimizing the energy flux for growth, we develop the equations governing the time evolution of cell morphology and growth rate in diverse environments. The resulting model accurately captures experimentally observed dependencies of bacterial cell size on growth rate, superlinear scaling of metabolic rate with cell size, and predicts nutrient-dependent trade-offs between energy expended for growth, division, and shape maintenance. By calibrating model parameters with available experimental data for the model organism *E. coli*, our model is capable of describing bacterial growth control in dynamic conditions, particularly during nutrient shifts and osmotic shocks. The model captures these perturbations with minimal added complexity and our unified approach predicts the driving factors behind a wide range of observed morphological and growth phenomena.

## I. INTRODUCTION

Living organisms face the challenge of continuously investing energy to fuel growth, biomechanical activities, and the maintenance of biomass. How energy uptake is allocated between the synthesis of new biomass and maintenance of existing mass to promote organism growth and reproduction is of central importance to ecology and evolution. Previous work has used energy budget models to understand ontogenetic growth in animals [1–5], where some fraction of the assimilated food is oxidized to sustain the total metabolic rate and the remaining fraction is synthesized and stored as biomass. While energy budget models have provided a quantitative and conceptual understanding of whole-organism growth in animals, how energy allocation strategies are implemented in single-celled microorganisms remains an open question. Here, we present a biophysical theoretical framework for dynamic allocation of cellular energy for key physiological tasks in bacteria, going beyond existing phenomenological models of microbial metabolism [6, 7]. Our theory quantifies how assimilated nutrient energy is dynamically allocated for bacterial growth, division, dissipation, and shape maintenance. We apply this framework to study bacterial growth and morphogenesis in steady and dynamic environments.

Much work has been done in recent years to understand how bacteria adapt their growth physiology in response to changes in nutrient environment [8]. Resource allocation models, based on coarse-grained partitioning of the cellular proteome into a few functional components, have provided a quantitative framework to predict how trade-offs between metabolic and gene expression machinery of the cell regulate bacterial growth rate [9–13]. While these theories are mostly focused on steady-state behaviors, recent studies have extended resource allocation models to describe transitions between steady states [14–18]. Despite these advances, existing resource allocation models are inadequate to describe physical behaviors of the cell, in particular how cellular resources are used for the control of cell shape, bioenergetics and mechanical activities. In addition, investigating perturbations from steady-state requires adding additional proteome sectors, which can quickly complicate the model [18]. On the other hand, mechanical models of cell growth have been developed to describe cell shape dynamics in growing bacterial cells [19, 20], but they do not take into account biochemical regulation and decision-making components of the cell. To circumvent these inadequacies, we introduce a new theoretical framework for bacterial growth and cell shape regulation based on allocation of uptake energy for physical and biochemical processes, including growth, division, storage and shape maintenance.

Through the optimization of energy flux for growth, we deduce the governing equations for cell size, shape and division dynamics. Upon calibrating model parameters using experimental data from the steady-state growth of the model organism *Escherichia coli*, we offer insights into how cellular energy uptake is distributed among various physiological tasks under specific nutrient conditions. Our theoretical framework aligns with multiple experimental observations, yielding predictions that shed light on the underlying driving forces of these phenomena. Our findings reveal a positive correlation between cell size and growth rate as a natural outcome of energy flux optimization. Analysis of cellular metabolic rates yields a superlinear scaling relationship between metabolism and bacterial cell size, explaining existing experimental data [21]. Furthermore, our model is adept at explaining complex transient behaviors observed in nutrient shift experiments [15]. Fitting our model to nutrient shift data [15, 22], we find that overshoots and undershoots past the final steady-state growth rate stem from disparate timescales governing nutrient importation and changes in cell morphology.

Notably, we showcase the model’s flexibility in capturing osmotic shocks [23] through manipulation of turgor pressure, highlighting its capability to predict energy reallocation under mechanical stress. Integrating the effects of both biochemical and mechanical regulation of growth, our unified approach offers a modular framework with broad applicability across organisms, offering a new perspective on the regulation of microbial growth physiology.

## II. DYNAMIC ENERGY ALLOCATION THEORY

### A. Energy flux balance and optimization principle

The starting point of our theory is the condition of energy flux balance, such that the rate of uptake of food energy, *E*_in_, is equal to the rate at which the consumed energy is used to fuel cellular metabolic processes (*E*_met_) plus the rate at which the accumulated energy is stored in cellular biomass (*E*_stored_) [5],

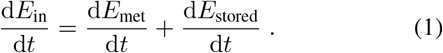

For a growing and replicating bacterial cell, we assume that the metabolic energy, *E*_met_, is used for four major tasks: (1) *E*_growth_, energy used for cellular growth and biosynthesis, (2) *E*_div_, energy expended for cell division, (3) *E*_mech_, mechanical energy expended for the maintenance of cell shape, structure and size, and (4) *E*_d_, the amount of energy lost due to dissipation. For simplicity, we consider a non-motile cell and neglect energy due to locomotion and other activities. A schematic depicting the energy partitioning scheme is shown in Fig. 1.

**FIG. 1.**
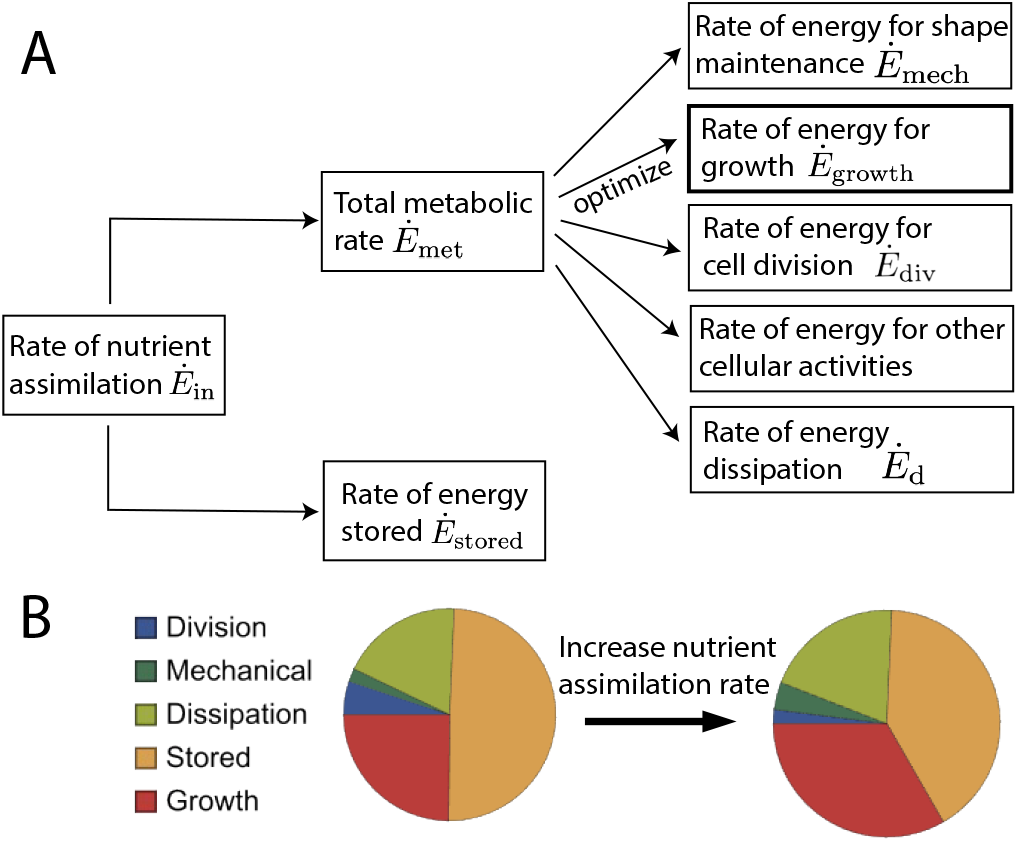
Dynamic energy allocation theory. (A) A schematic depicting how a bacterial cell partition assimilated nutrient energy into diverse cellular tasks. The energy utilized for growth, *Ė*_growth_, is highlighted to indicate that it is optimized to derive the governing equations of motion. (B) Energy allocation during cellular growth. The pie charts indicate fractions of assimilated energy *E*_in_ utilized for each energy sector at growth rate *κ* = 0.9 h^−1^ (left) and *κ* = 2 h^−1^ (right).

The energy components are functions of the state variables {*q*_*i*_(*t*) } that represent the the set of independent variables that describe the dynamic state of the cell. For a bacterial cell, these could be defined by cellular morphological parameters as well as protein copy numbers. While Eq. 1 is a general constraint for the cell, it is not sufficient to describe the full dynamics of {*q*_*i*_(*t*) } . To derive the equations of motion, we impose a principle of power optimization. In particular, for a growing bacterial cell, we hypothesize that the bacterium maximizes the rate of energy (or power) utilized for growth:

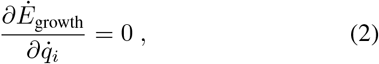

where dot denotes time derivative. This implies that the dynamics of the system are determined by the condition that the growth power, *Ė*_growth_, is a maximum with respect to 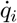, where 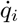 represents the rate of change in the current state *q*_*i*_(*t*).

Maximizing the rate of energy utilized for growth is similar to the maximum power principle often used in models of ecology and thermodynamics [24, 25]. In the case of bacteria, the rate of energy used for growth is proportional to the growth rate of the cell. The hypothesis for maximizing the growth power is motivated by experimental observations that *E. coli* cells evolve their metabolism towards a state that maximizes growth rate [26–28]. Furthermore, previous work has shown that growth-rate maximization leads to the empirical growth laws of bacterial cells [9–11, 29]. It is important to note that the maximum growth power hypothesis may not be applicable for bacteria under stress, such as under starvation, when cells may prioritize maintenance over growth [30].

### B. Modeling cellular energy components

Having described the central theoretical concepts and model hypothesis, we now turn to modeling the different components of cellular energy.

#### Nutrient intake

Assuming that the nutrient uptake occurs uniformly through the cell surface, the energy for nutrient intake, *E*_in_, is taken to be proportional to the surface area *S* of the cell, such that *E*_in_ = *εS*, where *ε* is proportional to the surface concentration of nutrients as well as the surface density of transporters.

#### Stored energy

The stored energy in the cell is given by *E*_stored_ = *gV*, where *V* is the cell volume and *g* is the energy stored per unit volume of the biomass.

#### Turgor pressure

The energy due to turgor pressure is given by *E*_turgor_ = − *PV*, where *P* is the turgor pressure and *V* is the cell volume.

#### Cell shape maintenance

The mechanical energy for maintaining the shape of the cell envelope, *E*_mech_, is given by the sum of the strain energy in the bacterial cell envelope (*E*_strain_), and the bending energy required to maintain cell curvature (*E*_curv_):

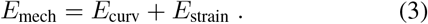

For a rod-shaped bacterial cell of length *L* and radius of cross-section *R*, the curvature energy is given by

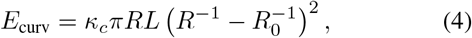

where *R*_0_ is the preferred radius of cross-section and *κ*_*c*_ is the bending rigidity. To compute the strain energy in the cell envelope due to turgor pressure *P*, we assume that the rate of cell growth is slower than mechanical relaxation, such that the cell envelope is always in mechanical equilibrium. The strain energy in the peptidoglycan cell wall can be expressed as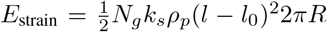, where *N*_*g*_ is the number of glycan strands, *ρ*_*p*_ is the density of peptide crosslinkers, *k*_*s*_ is the stifness of a crosslinker, *l* is the deformed length and *l*_0_ is the undeformed (rest) length of the crosslinker. In mechanical equilibrium, stress in the cell wall balances the turgor pressure, such that the length deformation is given by: *l* −*l*_0_ = *PR/*(2*k*_*s*_*ρ*_*p*_). As a result, we can rewrite the strain energy as

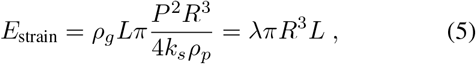

where *λ* = *P* ^2^*ρ*_*g*_*/*4*k*_*s*_*ρ*_*p*_ can be interpreted as the mechanical stress per unit circumference, and *ρ*_*g*_ = *N*_*g*_*/L* is the line density of the glycan strands (assumed to be uniform). We note that the scaling form for the strain energy in (5) agrees with previous calculations done using thin shell elasticity theory for a rod-shaped bacterial cell [19].

#### Cell division

Bacterial cell division relies on a large protein complex called the divisome, which orchestrates the assembly of the Z-ring around the mid-cell region [31]. The Z-ring is composed of FtsZ filaments, capable of generating constrictive bending forces [32, 33]. Moreover, the divisome triggers peptidoglycan synthesis and guides the formation of the septum [34]. Previous work has demonstrated that FtsZ generates a small amount of mechanical force that is insufficient to fully constrict the mid-cell [20, 35, 36]. Instead, the primary driving force for cell constriction arises from cell wall synthesis at the septum [20, 37, 38]. Consequently, we disregard the mechanical energy cost of Z-ring constriction and model the energy of cell division as being driven by the synthesis of division proteins (e.g., FtsZ).

Defining *X* as the total abundance of division proteins, the energy for the synthesis of division proteins can be written as

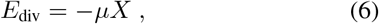

where *μ* is the chemical potential for division protein synthesis. In most bacterial cells that follow the adder mechanism for cell division control [8, 39], cell division is triggered once a threshold amount of division proteins are accumulated during the cell cycle [22, 40–42]. Since FtsZ abundance in the Z-ring reaches a fixed threshold prior to division [41], assume that the threshold abundance of division proteins, *X*_0_, is proportional to the cell diameter: *X*_0_ = 2*πRγ* [43], where *γ* is the proportionality constant, and *R* is the cell radius. Therefore, the energetic cost of division during a cell cycle is ≈ −2*πμγR*.

*Energy dissipation –* Living cells can be considered as open thermodynamic systems that dissipate energy as they grow and replicate [44]. For a growing bacterial cell, the rate of energy dissipation (d*E*_d_*/*d*t*) is a sum of energy dissipation rate due to mechanical (D_mech_) and chemical components (D_chem_):

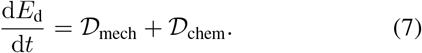

In non-equilibrium thermodynamics, a commonly made assumption is that the energy dissipation rate follows a quadratic relationship with the “rate” of change in state variables, known as the linear response theory [45–47]. Assuming linear response theory, the dissipation rate of mechanical energy can be taken to be proportional to the square of the strain rate. Specifically, for a rod-shaped bacterial cell with length *L* and radius *R*, the strain rates can be expressed as 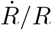 and 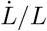. Consequently, we can formulate the mechanical dissipation rate as:

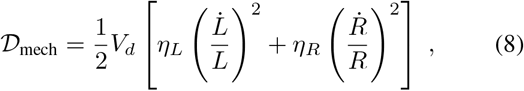

where *η*_*L*_ and *η*_*R*_ are the viscosity parameters and *V*_*d*_ is the volume over which dissipation occurs. We take *V*_*d*_ = *hS*, where *h* is the cell envelope thickness, and *S* ≈ 2*πRL* is the surface area of dissipation for a rod-shaped cell. Analogously, the dissipation rate of chemical variables of the system can be assumed to follow a quadratic relationship with the square of the rate of production of that chemical.

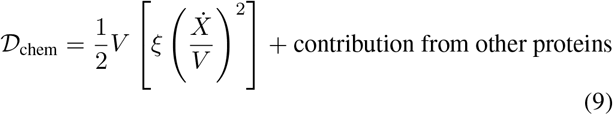

where *ξ* is a constant. Therefore, the total dissipation rate due to cell envelope growth and division is:

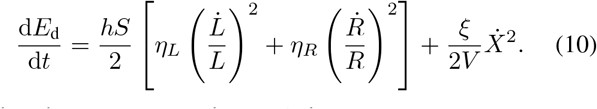

The above form for the total dissipation rate is consistent with thermodynamic principles since the resulting dissipative forces, 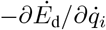, are not symmetric under time-reversal.

*Energy for growth –* Cellular growth energy involves the energy required to synthesize biomass and the surface area. Here we do not directly define the growth energy, but instead calculate it from the energy flux balance relation in Eq. (1). We can write the flux of energy used for growth as:

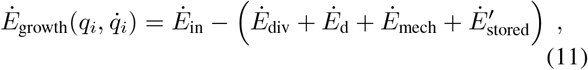

where 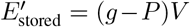 is the effective stored energy. In exponentially growing cells, *Ė*_growth_ is proportional to the growth rate *κ*, which follows from the equations derived in the next section.

### C. Equations governing cell growth and shape dynamics

For a rod-shaped bacterial cell such as *E. coli* or *B. subtilis*, the minimal set of state variables required to describe its growth and division dynamics are the cell length *L*, radius *R* and division protein abundance *X*. Thus, *q* ={ *L, R, X* } . Using Eq. 2, the condition for optimizing growth power, we obtain

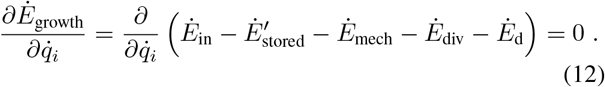

Since *E*_in_, *E*_stored_, *E*_div_, and *E*_mech_ are functions of *q*_*i*_’s only, then 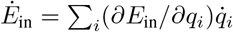, and similarly for the rest. Therefore the governing equations for the dynamics of *q*_*i*_ are given by:

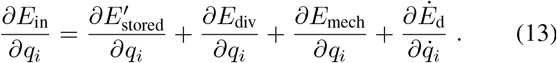

As a first step, by optimizing *Ė*_growth_ with respect to 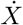, we can obtain:

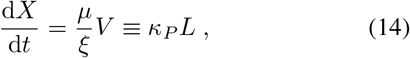

where we defined *κ*_*P*_ = *πR*^2^*μ/ξ*, as the rate of division protein synthesis per unit length. Eq. (14) is identical to the model for volume-specific production of division proteins as considered previously by several authors [13, 22, 40–42]. Using Eqs (2) and (11), we derive the equations governing cell length and radius:

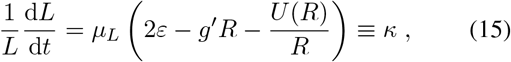

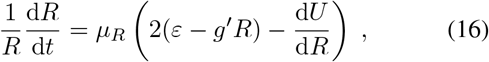

where *μ*_*L*_ = 1*/*2*η*_*L*_*h, μ*_*R*_ = 1*/*2*η*_*R*_*h, U* (*R*) = *E*_mech_(*R, L*)*/πL, g*^t^ = *g* − *P*, and *κ* is the longitudinal growth rate. From (16) we see that cells maintain a steady-state radius in a given growth condition to minimize an effective potential *E*_eff_ defined as

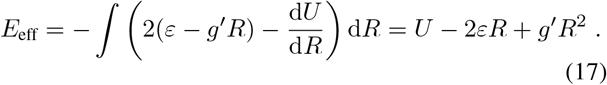

With a fixed radius at steady-state, cell length increases exponentially as expected for rod-shaped bacteria. Exponential growth is contingent on sufficient energy intake (per unit surface area) *ε* such that *κ >* 0. In other words, the energy flux into the cell must surpass the mechanical energy costs and anticipated energy storage for the cell to grow. As seen in Fig. 2A, in nutrient-poor conditions, cells would need to decrease their stored energy density *g* (from the value we will later extract from experimental data) to permit *κ >* 0. Using Eqs. (14)-(16) we can simulate the dynamics of length, radius and protein abundance over multiple generations of growth and division as shown in Fig. 2B-C. Division occurs once *X* reaches a fixed threshold, as discussed earlier. Upon changes in growth rate, induced by changes in *ε*, the landscape for the effective potential changes such that cells find a new radius that minimizes *E*_eff_ as seen in Fig. 2D.

**FIG. 2.**
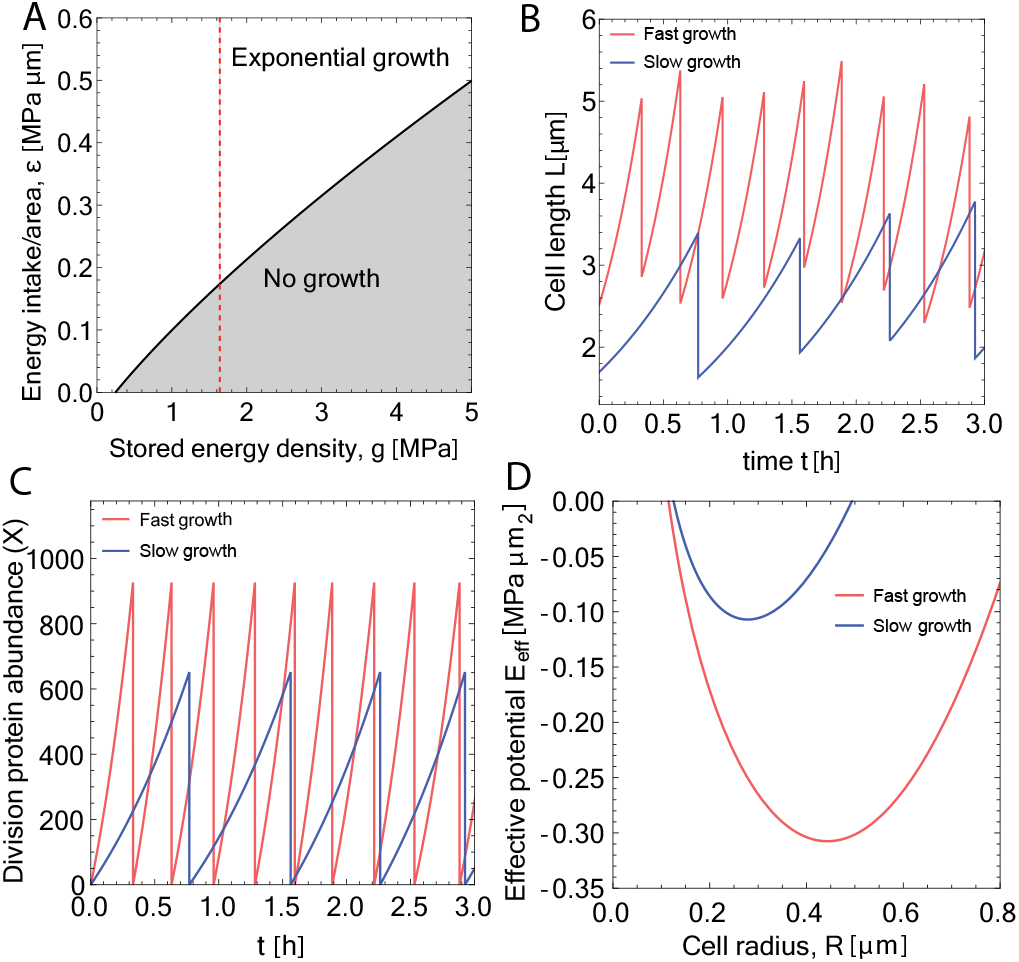
Single-cell growth dynamics in the energy allocation model. (A) Phase diagram showing regions of parameter space permitting exponential growth. (B) Representative trajectories of cell length (using Eq. 15) for *κ* = 0.9 h^−1^ (blue) and *κ* = 2 h^−1^ (red). Noise is included in the initial size of daughter cells and cell division ratio to illustrate model stability. (C) Representative trajectories of division protein abundance (using Eq. 14) for *κ* = 0.9 h^−1^ (blue) and *κ* = 2 h^−1^ (red). (D) Effective potential energy landscapes (Eq. 17) in nutrient-poor (*κ* = 0.9 h^−1^, blue) and nutrient-rich environments (*κ* = 2 h^−1^, red). The minimum value of each curve corresponds to the steady-state radius.

Before constraining the model parameters from experimental data, we can obtain some general insights about the interdependence between cell size and growth. Increasing the nutrient influx *ε* increases the growth rate *κ* (Eq. 15, Fig. 2B), as well as the division rate *κ*_*P*_ (Fig. 2C). As *ε* increases, we see that the steady-state value for cell radius *R* must increase to compensate (Fig. 2D). Conversely, an increase in energy storage *g* (while keeping *ε* constant) prompts a reduction in *R*. In simpler terms, a greater nutrient influx is associated with increased cell width, while greater energy storage is linked to decreased width. As evident from Eq. (15), the growth rate decreases with increasing mechanical energy. With an increase in either the bending stifness *κ*_*c*_ or the cell envelope stress *λ*, cell width would decrease as the energy cost of maintaining a larger envelope is greater. Thus, without specifying the numerical values for the parameters, cell width naturally increases with the growth rate in our model, consistent with experimental data [48].

Unlike the control of cell width, the average cell length is set by the division protein accumulation threshold, *X*_0_, and the division protein production rate, *κ*_*P*_ . Integrating Eqs. (15) and (14) across a cell cycle, we arrive at Δ*L* = *X*_0_*κ/κ*_*P*_, where Δ*L* is the length added over a cell cycle. Thus the added length Δ*L* increases with *κ*. Δ*L* also increases as *X*_0_ increases since the cell needs to grow for longer to reach the threshold. Conversely, the increase in the production rate *κ*_*P*_ leads to a decrease in Δ*L*.

### D. Parameter determination

With the theoretical framework in place, we now turn to determining the model parameters using available experimental data on the model organism *E. coli*. All parameters discussed in this section are listed in Table I for ease of reference. These include mechanical, geometrical and biochemical parameters of the cell. Fig. 3A documents our procedure of parameter determination. Several mechanical parameters such as the bending rigidity *κ*_*c*_, cell-wall stifness *k*_*s*_ and turgor pressure *P* can be fixed from reported measurements. From prior work, we estimate *κ*_*c*_ = 0.03 MPa *μ*m^3^ [35], and assume that the preferred radius of cross-section *R*_0_ is the same as the preferred radius of curvature for MreB (200-400 nm [49]) which hold the circumferential structure of the cell wall. The parameter *λ* = *P* ^2^*ρ*_*g*_*/*4*k*_*s*_*ρ*_*P*_ quantifies the mechanical stress on the cell envelope, which can be indirectly estimated. Prior work on thicker gram-positive cell walls has shown that *k*_*s*_*ρ*_*p*_ ≈ 127 MPa [51] for each layer, of which the gram-negative *E. coli* has only one. The turgor pressure in *E. coli* cells is *P* = 0.3 MPa [52] and *ρ*_*g*_ = 1000 *μ*m^−1^ [51], leading to *λ* ≈ 0.18 MPa/*μ*m.

**TABLE I.** List of model parameters.

| Parameter | Value | Physical meaning | Source |
| --- | --- | --- | --- |
| $\kappa_0$ | varied | steady-state growth rate | Initial condition |
| $R_0$ | $0.3\mu\text{m}$ | MreB radius of curvature | [49] |
| $\kappa_c$ | $0.03\text{ MPa } \mu\text{m}^3$ | bending rigidity | [20] |
| $k_s\rho_p$ | $127\text{ MPa}$ | peptide crosslinker stiffness | [50] |
| $P$ | $0.3\text{ MPa}$ | turgor pressure | [20] |
| $\rho_g$ | $1000\text{ } \mu\text{m}^{-1}$ | glycan line density | [51] |
| $h$ | $5 \cdot 10^{-3}\mu\text{m}$ | envelope thickness | [20] |
| $\gamma$ | $364\text{ FtsZ}/\mu\text{m}$ | division protein threshold/length | [50] |
| $R^*$ | $(0.1h\kappa_0 + 0.195)\mu\text{m}$ | steady-state radius | [39] |
| $L_0$ | $1.26\exp(0.33h\kappa_0)\mu\text{m}$ | newborn length | [39] |
| $\mu$ | $9.2 \cdot 10^{-5}\text{ MPa } \mu\text{m}^3$ | chemical potential for division protein synthesis | Estimated |
| $\lambda$ | $0.18\text{ MPa}/\mu\text{m}$ | envelope stress | Derived |
| $\eta_L$ | $1/2h\mu_L$ | viscosity (length) | Definition |
| $\eta_R$ | $1/2h\mu_R$ | viscosity (radial) | Definition |
| $V_d$ | $h\text{ S}$ | dissipation volume | Definition |
| $\xi$ | $\mu/\kappa_p$ | dissipation rate for protein production | Derived |
| $\kappa_p$ | (20) | division protein production rate | Initial Condition |
| $\mu_L$ | (19) | longitudinal mobility | Initial Condition |
| $\varepsilon$ | (18) | average energy import | Initial Condition |
| $\mu_R$ | $[10^{-1}, 10^2]\text{ MPa}^{-1}\text{h}^{-1}\mu\text{m}^{-1}$ | radial mobility coefficient | Fit |
| $g$ | $1.64\text{ MPa}$ | stored energy density | Fit |

**FIG. 3.**
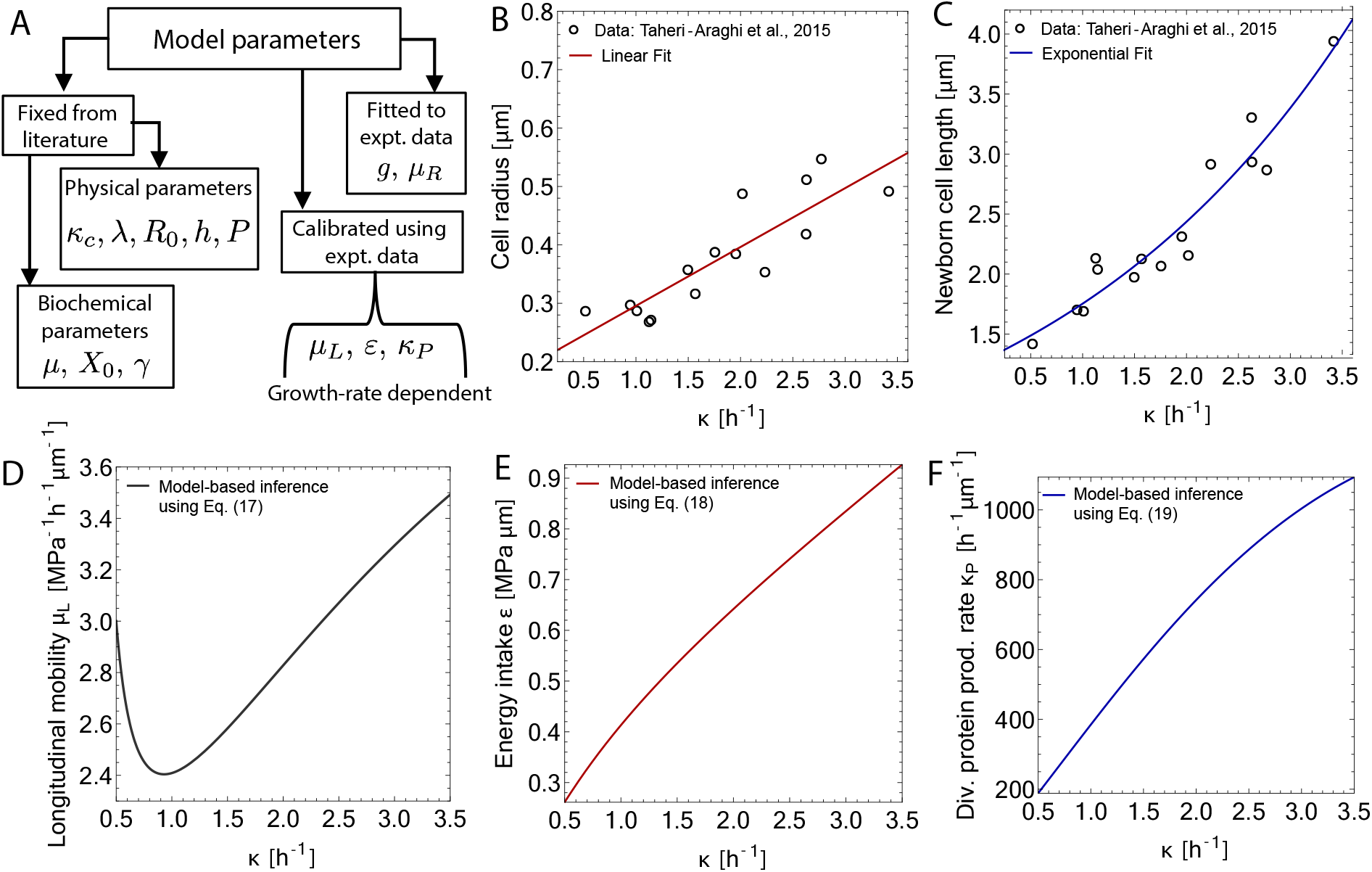
Calibration and inference of model parameters. (A) A schematic illustrating how the model parameters are determined. (B) Dependence of steady-state cell radius on the nutrient-specific growth rate *κ* for exponentially growing *E. coli* in a variety of different nutrient conditions [39]. We fit the experimental data (open circles) using a linear model *R*(*κ*) = (0.1h *κ* + 0.195)*μ*m (solid line). (C) Measured values of newborn cell length *L*_0_ of exponentially growing *E. coli* in a variety of different nutrient conditions [39]. We fit the dependence of *L*_0_ on growth rate using an exponential model *L*_0_(*κ*) = 1.26 *μ*m exp(*κ* 0.33 h). (D) Model-based inference of longitudinal mobility of the cell, *μ*_*L*_, as a function of growth rate, determined using Eq. (19). (E) Model-based inference of energy intake per unit area, *ε*, as a function of growth rate, determined using Eq. (18). (F) Model-based inference of the division protein production rate *κ*_*p*_ as a function of growth rate, determined using Eq. (20).

The chemical potential for division protein production can also be estimated from available data. Given that FtsZ contains 383 amino acids and consumes 4 ATP per added amino acid to a peptide chain [53], coupled with an energy output of 36; *kJ* per mole of ATP, we deduce *μ* = 9.2 10^−5^ MPa *μ*m^3^/FtsZ. Furthermore, there are approximately 3200 FtsZ molecules per cell [50]. Considering that 30% of the total FtsZ is present in the ring [54], we can estimate the thresh-old abundance of FtsZ to be *X*_0_ ≈ 960. For *E. coli* cells with radius *R* = 0.42*μ*m, we arrive at an approximate value of FtsZ abundance per unit circumference *γ* ≈ 364 FtsZ*/μ*m.

The model comes with several other unknown parameters that may be dependent on the specific growth conditions. These include *ε* (energy intake per unit area), *g* (stored energy density), *μ*_*L*_ (longitudinal mobility), *μ*_*R*_ (radial mobility) and *κ*_*p*_ (division protein synthesis rate). We can determine these parameters from given experimental conditions. The first condition is that the radius measured under steady-state exponential growth [39], *R*^∗^, corresponds to the radius *R* that minimizes the effective potential in Eq. (17). Using this condition, we obtain the energy intake per unit area *ε*:

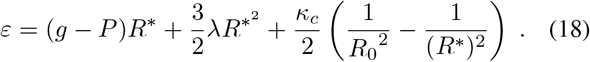

From experimental data on the steady-state exponential growth of bacterial cells [39], we have the second experimental condition that *L*^−1^*dL/dt* = *κ*_0_, where *κ*_0_ is the nutrientspecific growth rate. Taken together with Eq. 15, we get

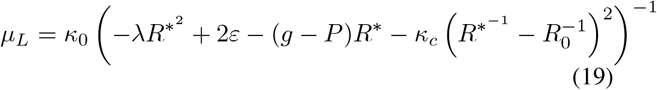

To find the division protein synthesis rate *κ*_*p*_, we integrate Eq. (14) and (15) with the boundary condition that *X* = *X*_0_ = 2*πRγ* at the time of cell division. This yields,

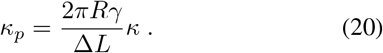

Under fixed aspect ratio of *E. coli* cells [43], the above equation reduces to the condition of balanced biosynthesis *κ*_*p*_ ∝ *κ* [41], i.e. division protein production rate is proportional to the growth rate. Fits to experimental data for how steadystate cell radius (*R*^∗^) and newborn length (*L*_0_) change with growth rate is provided in Figs. 3B-C, which make predictions for how the parameter *μ*_*L*_, *ε*, and *κ*_*p*_ change with growth rate (Figs. 3D-F).

The remaining two parameters *μ*_*R*_ (radial mobility coefficient) and *g* (stored energy density) are determined by fitting the model to experimental data on nutrient shifts (see Section IV). *μ*_*R*_ controls how fast the cell radius adapts to a new steady-state value when perturbed, but it does not influence the cell morphology or energies during steady-state exponential growth. The value of *μ*_*R*_ depends on the type of perturbation, as seen later in sections IV and V. While *g* also does not affect steady-state dynamics, it does affect the steady-state values of *E*_stored_, and *E*_growth_. In section IV we discuss how *g* is extracted from nutrient downshift data. Note that, unlike *μ*_*R*_, *g* does not change with *κ* and thus does not depend on the nutrient-specific growth rate.

## III. ENERGY ALLOCATION STRATEGIES AND METABOLIC SCALING LAW

### A. Trade-offs in energy allocation

After determining the model parameters in various growth conditions, we employ the model to predict the evolution of cellular energies throughout the cell cycle and their dependence on growth rate. As cells elongate, energy in each sector accumulates in proportion to cell size (refer to section II B and II C for definitions). Consequently, the energy of each sector *E*_*i*_(*t*) increases over the course of the cell cycle (Fig. 4A) and is then split equally amongst the daughter cells at division. The rate at which energy accumulates in each sector, **Ė**_*i*_, is directly proportional to *L*, as the energies themselves are proportional to *L*, thereby ensuring that the allocation to different sectors remains independent of the cell cycle. In other words, our model predicts no shifts in energy densities (*E*_*i*_(*t*)*/L*(*t*)) in exponentially growing cells at steady-state.

**FIG. 4.**
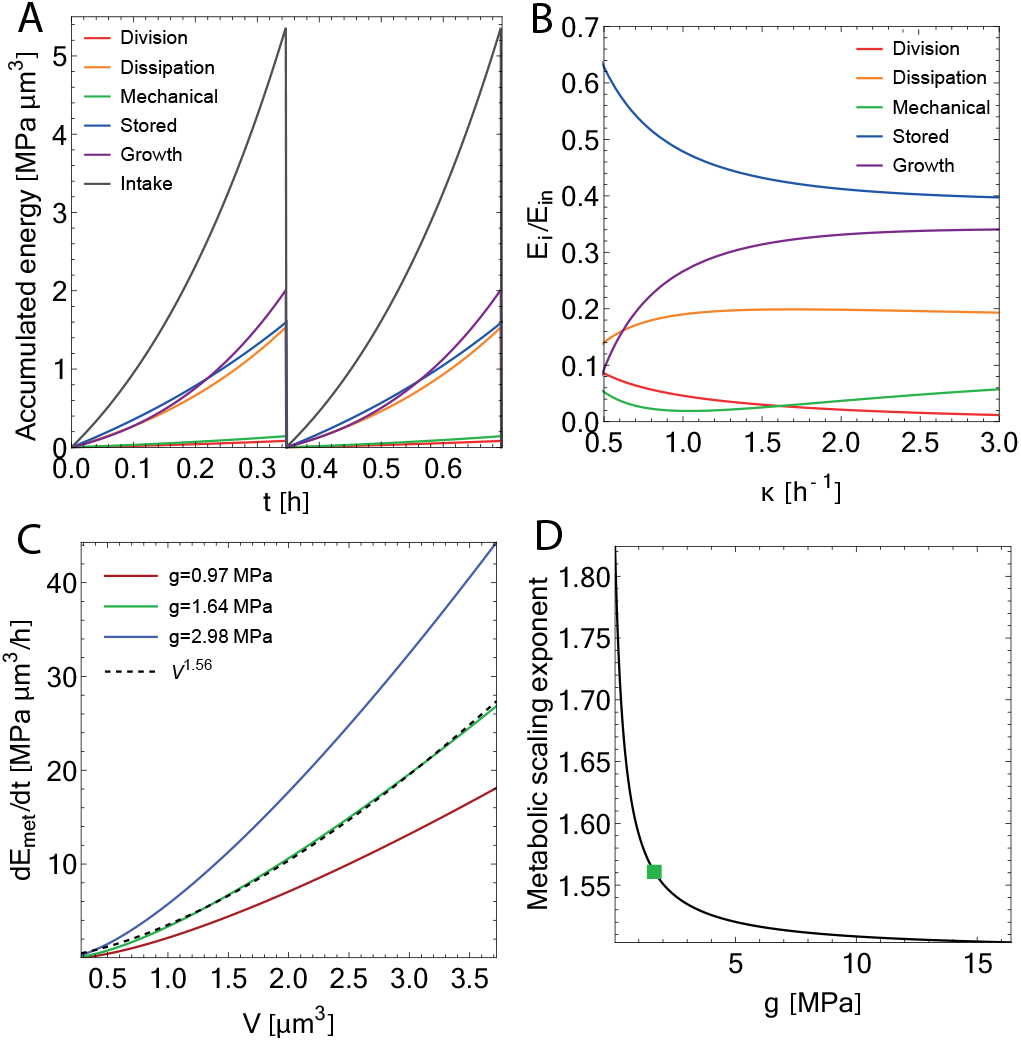
Energy allocation trade-offs and metabolic scaling law. (A) Accumulated energy since birth as a function of time, through two cycles of growth at *κ* = 2 h^−1^. The intake energy (black line) is split amongst the various energy sectors, all of which grow at a rate proportional to the *L*. (B) Energy in each sector, *E*_*i*_, normalized by the total intake energy, as a function of growth rate. *E*_growth_ is calculated by subtracting all other energy sectors from *E*_in_. (C) Metabolic rate as a function of cell volume, as defined in Eq. (1). Three curves are shown for different values of the stored energy density *g*, with the green curve (*g* = 1.64 MPa) representing the scaling law observed experimentally (see section IV). The scaling exponent is fit to be approximately 1.56 for this value of *g*. Smaller values of *g* (red line) have a lower scaling exponent while larger *g* (blue line) results in higher scaling exponents. (D) Metabolic scaling exponent as a function of *g*. The scaling exponent 1.56 for the calibrated value of *g* = 1.64 MPa is marked with a solid green square.

In terms of the absolute values, all energies increase as a function of growth rate *κ*, since cell length and radius both increase with *κ*. However, normalizing cell-cycle averaged accumulated energy by the intake energy is more insightful, as it indicates how intake energy is allocated in different sectors (Fig. 4B). We thus define energy fractions relative to the intake energy as

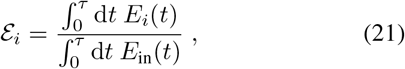

where *i* ∈ {‘growth’, ‘stored’, ‘mech’,’div’,’d’} and *τ* is the cell cycle duration. As *κ* increases, we see a minimum of ε_mech_ at growth rate for cell radius *R* = *R*_0_, with greater energy costs the further the cell deviates from that optimum growth rate (Fig. 4B). ε_div_ decreases with growth rate, as *E*_div_*/E*_in_ decreases with cell size (∝ 1*/V*, Fig. 4B). Notably, cells waste more energy through dissipation (ε _d_) during faster growth, given the nonlinear scaling of *E*_d_ with cell size. One of the more drastic changes is a decrease in ε _stored_ with growth rate (Fig. 4B), as cells utilize a greater fraction of the overall intake energy during fast growth conditions. Together, the changes in these sectors result in an increase in the growth energy fraction, ε _growth_, when cells are exposed to more optimal growth conditions. These predicted changes in energy fractions with growth rate are reminiscent of changes in proteome fractions with growth rate [10, 11, 55, 56]. The proteome fraction of the proteins regulating cell growth, such as ribosomal proteins, increases with growth rate similar to how the energy fraction allocated to growth increases with growth rate.

### B. Scaling of metabolic rate with cell size

The scaling of metabolic rate with organism size represents a fundamental characteristic trait of living organisms. It has been shown previously that the metabolic rate of birds and mammals follows a scaling law of 3/4th power relative to body size [57]. In contrast, microorganisms like bacteria display super-linear scaling of metabolic rate with cell size [21]. The biophysical origin of this super-linear scaling remains a mystery and to date, there is no mechanistic model explaining this scaling relation. Our theory now enables us to directly compute the metabolic rate, **Ė**_met_, and investigate its scaling relations with cell size under various parameter values. Crucially, as elaborated below, our findings indicate that the super-linear scaling of metabolic rate with cell size naturally emerges from the increase in bacterial cell size with growth rate.

Using the equation for metabolic rate as defined in Eq. (1), and substituting our expressions for *E*_in_ and *E*_stored_, we have

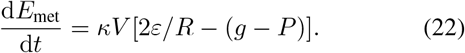

If *ε* and *g* were both independent of growth rate, one might expect the metabolic rate to decrease with cell size, since the positive term scales like *R*^−1^. However, *ε* is growth rate-dependent and thus increases with cell size. Substituting the expression for *ε* in Eq. (18) into Eq. (22) we get:

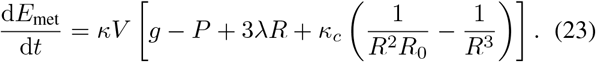

Using this expression, we see that the metabolic rate scales approximately as a power of cell volume (Fig. 4C). Since *V* (*κ*) is concave (as *L*(*κ*) is exponential, Fig. 3Bf-C), *κ*(*V*) is sublinear and contributes to the overall superlinear scaling. Conceptually, the parameter *g* plays a role in determining this scaling as it controls both *E*_stored_ and *ε* (see Eq. 18). As shown in Fig. 4D, smaller values of *g* correspond to higher scaling exponents and larger values of *g* correspond to smaller scaling exponents. In other words, a greater amount of energy stored in the cellular biomass naturally corresponds to a lower metabolic rate. Analytically, we see that if *g* dominates Eq. (23), then the scaling exponent is less influenced by the terms that have *R* dependence. At *R* = *R*_0_, the *g, λ, κ*_*c*_ terms in (23) are approximately 1.64 MPa, 0.16 MPa, and 0 MPa, respectively. Even when *R* deviates from *R*_0_, we see that *g* is an order of magnitude larger than the second largest term.

Regardless of the parameter choices, our model always predicts superlinear scaling of metabolic rate with cell size for bacteria. This is in contrast to animals [5] and other microorganisms [21], which typically exhibit sublinear scaling. The key distinction that sets bacteria apart is that growth rate increases with cell size. Independence of growth rate and cell size would result in nearly linear scaling as *g* dominates the terms (23), whereas a decreasing cell size with growth rate would instead result in sublinear scaling of metabolic rate with cell size.

## IV. DYNAMIC RESPONSE TO NUTRIENT SHIFTS

With an understanding of cellular energy allocation strategies during steady-state exponential growth, we turn to modeling cellular response to perturbations about steady-state. To this end, we first inquired whether our model is capable of predicting growth rate dynamics in response to rapid shifts in nutrient quality in the growth medium. The change in nutrient quality can be simulated by dynamically changing the parameters *ε* (Eq. 18) and *μ*_*L*_ (Eq. 19), since they are dependent on the nutrient-specific growth rate (see Fig. 3D-E). As a simple way to represent a gradual time-dependent response, we model nutrient-induced changes in *ε* and *μ*_*L*_ as logistic functions,

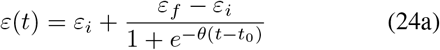

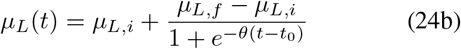

where the subscript *i* corresponds to the initial value of the parameter and *f* corresponds to the final value, *θ* is the steepness of the function, and *t*_0_ is the time when the shift is applied. As indicated in Eqs. (15) and (16), the changes in *ε* and *μ*_*L*_ gradually bring the growth rate and cell size to the new steady state. The parameter *μ*_*R*_ introduced in Eq.(16) governs the rate at which the cell radius reaches the new equilibrium value after the shift, while the undetermined parameter *g* controls the timescale over which the growth rate attains the new steadystate value. Consequently, for a given nutrient shift, we have a total of four undetermined parameters: *θ, t*_0_, *μ*_*R*_, and *g*, all of which are determined through fitting the model predictions for the growth rate to experimental data [15].

Fig. 5 shows model predictions for cell size and growth rate dynamics during nutrient upshifts (Fig. 5A-D) and downshifts (Fig. 5E-H). The model is fitted to experimental data from Ref. [15], and the comparisons for growth rate dynamics during nutrient upshift and downshift are presented in Figs. 5A and 5E, with corresponding predictions for cell radius *R*(*t*) and energy intake *ε*(*t*) shown in Figs. 5B,F. During nutrient upshift, the growth rate changes gradually (Fig. 5A), and *R*(*t*) closely follows the trajectory of *ε*(*t*) (Fig. 5B). In contrast, during a nutrient downshift, the growth rate initially decreases sharply, undershoots the new steady-state, and gradually rebounds to a new steady-state (Fig. 5E). Unlike the upshift, *ε*(*t*) quickly adjusts to its new value, while *R* takes hours to reach the new steady-state (Fig. 5F). Our model suggests that the difference between upshift and downshift dynamics is influenced by the timescale at which nutrient import (*ε*) stabilizes to the new steady-state and whether or not there is a lag between *ε* and cell radius *R*. In a nutrient downshift, nutrient removal (*ε*) occurs much faster than growth, resulting in a temporal lag between *ε* and *R* (Fig. 5F). Conversely, during an upshift, while the new nutrients become available externally at a similar timescale to the downshift, cells may need to upregulate nutrient import and protein synthesis to fully utilize their richer environment. These processes are inherently connected to cell growth and cell envelope synthesis, leading to no observed lag in *ε* and *R* (Fig. 5B).

**FIG. 5.**
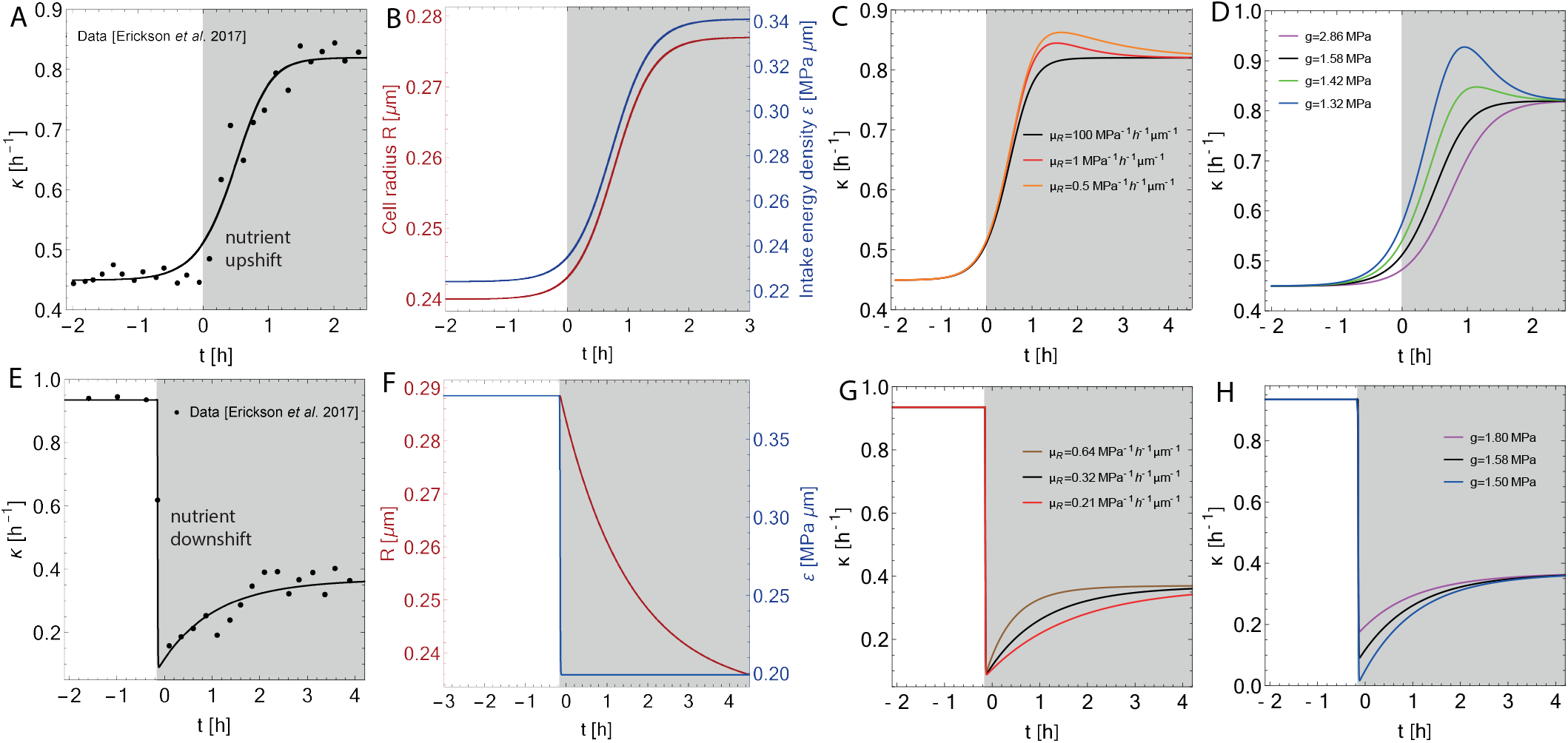
Modeling response to nutrient upshifts and downshifts. (A) Growth rate vs time for a representative nutrient upshift experiment, with the experimental data shown in black solid circles and model fit shown in solid black line. The sample experimental data, taken from Ref. [15], is for the *E. coli* strain NCM3722, grown in 20 mM succinate with a co-utilized substrate of 0.2% glucose added/removed at *t* = 0 for the upshift/downshift. Growth rate *κ* is calculated by integrating Eq. (15). The fitting parameters are *θ* = 3.1 h^−1^, *μ*_*R*_ = 100 MPa^−1^h^−1^*μ*m^−1^, and *g* = 1.64 MPa. The white background indicates pre-shift and the grey region indicates post-shift environment. (B) Predicted trajectory of cell radius *R*, and energy intake per unit area, *ε*, as functions of time for the data shown in panel (A). (C) Predicted growth rate dynamics during nutrient upshift for different values of *μ*_*R*_. (D) Predicted growth rate dynamics during nutrient upshift for different values of *g*. (E) Growth rate vs time for a representative nutrient downshift experiment, with the experimental data shown in black solid circles and model fit shown in solid black line. The fitting parameters are *θ ≈* 10^5^ h^−1^ (effectively a step function in *ε*), *μ*_*R*_ = 0.32MPa^−1^h^−1^*μ*m^−1^, and *g* = 1.58 MPa. (F) Predicted time evolution of *R* and *ε* for the nutrient downshift data shown in panel (E). (G) Predicted growth rate dynamics during nutrient upshift for different values of *μ*_*R*_. (H) Predicted growth rate dynamics during nutrient upshift for different values of *g*.

To examine how changes in model parameters impact the response to nutrient shifts, we simulated single nutrient upshifts and downshifts with varying values of *μ*_*R*_ (Fig. 5 C/G) and *g* (Fig. 5D/H). In upshift simulations, reducing *μ*_*R*_ from the fitted value introduces a lag between *ε*(*t*) and *R*(*t*), causing *ε*(*t*) to reach the new steady-state value before *R*(*t*). This results in a trajectory with an overshoot in growth rate and an increased time needed to settle into the new steady-state (Fig. 5 C). Similarly, decreasing *g* from the fitted value also leads to a growth rate overshoot (Fig. 5D), while increasing *g* does not qualitatively alter the growth rate trajectory and is essentially equivalent to delaying the time of the upshift. We note that growth rate overshoots have been reported recently in microfluidic nutrient upshift experiments [22].

In nutrient downshift simulations, increasing (decreasing) *μ*_*R*_ decreases (increases) the time required for the growth rate to attain the new steady state, without altering the amplitude of the undershoot (Fig. 5G). Conversely, smaller values of *g* result in a more pronounced undershoot in the growth rate, while the time to reach the new steady state remains unaffected. We observe that *g* is the sole parameter governing the depth of the growth rate undershoot in response to nutrient downshifts, enabling us to deduce its value through model fitting to data. Utilizing multiple nutrient downshift datasets from Ref [15], we do not observe a dependence of the fitted value of *g* on the initial and final growth rates. Fitting three datasets yields *g* = 1.58 MPa, *g* = 1.61 MPa, and *g* = 1.73 MPa, which are not correlated with the magnitude of the downshift or initial growth rate in the available data. This is not unexpected as *g* is a stored energy density rather than the absolute stored energy which is influenced by initial and final growth rate dependant cell size). Consequently, we used the average fitted value of *g* (≈ 1.64 MPa) for the majority of simulations in this paper.

As growth rate and cell size change in response to a nutrient upshift, the cell’s energy allocation strategies follow the anticipated growth rate dependencies, as shown in Fig. 4. However, this pattern does not hold for a downshift due to the disparate adaptation timescales of *R* and *ε*. The *ε* dependence is solely present in *E*_in_, resulting in a rapid reduction in intake energy at *t* = 0, while the other energy sectors remain unaffected as *R* has not yet changed. Consequently, *E*_div_, *E*_mech_, *E*_stored_, and *E*_d_ temporarily account for a larger fraction of *E*_in_. As a result, there is a diminished fraction of *E*_in_ available for growth compared to the steady-state scenario, leading to the observed undershoot.

## V. ADAPTATION TO OSMOTIC SHOCKS

An advantage of the energy allocation theory is that it provides a unified framework to study both mechanical and biochemical perturbations. Here we employ the model equations to study cellular growth response to osmotic shocks. Hyperosmotic shocks, characterized by increases in external osmolarity, result in an immediate reduction in cytoplasmic water content, leading to a decrease in growth rate and eventual plasmolysis for sufficiently prolonged shocks [23]. Conversely, hypoosmotic shocks, involving a decrease in external osmolarity, lead to an immediate increase in cytoplasmic water content and subsequent adaptation, with cells ultimately relaxing their growth rate to a sustainable value closer to the prior steady-state [23, 58]. Changes in external osmolarity directly impact the turgor pressure term in the energy, consequently changing the strain energy *E*_strain_ and the resulting equations of motion. Therefore, given that pressure changes initially occur on a timescale smaller than cell growth, we begin by examining the effects of rapid changes in turgor pressure on cellular growth and shape.

### A. Hyperosmotic shocks

We first consider the case of a hyperosmotic shock, which is modeled as a step-function decrease in turgor pressure from its initial value *P*_0_ to a final value *P*_*f*_ (Fig. 6A). This results in a sharp decline in the growth rate, followed by a gradual recovery to a new steady-state value below the pre-shock level (Fig. 6B). The extent of recovery depends on the shock magnitude; if the shock is mild, the growth rate fully returns to the pre-shock value (Fig. 6A-B). The reduced pressure also drives a gradual reduction in cell size, with both cell length and radius decreasing to a new steady-state value post-shock (Fig. 6C-D). The reduction in cell radius facilitates growth rate recovery over time, given the negative feedback between *κ* and *R* as evident from Eq 15. Moreover, the decreased radius reduces the length increment during the cell cycle, as the threshold amount of division proteins *X*_0_ is proportional to the cell radius. However, this reduced cell radius contradicts experimental data [59], where it is observed that the cell radius eventually returns to its pre-shock value. Additionally, the growth rate is observed to return to its pre-shock value even for substantial shock magnitudes [23].

**FIG. 6.**
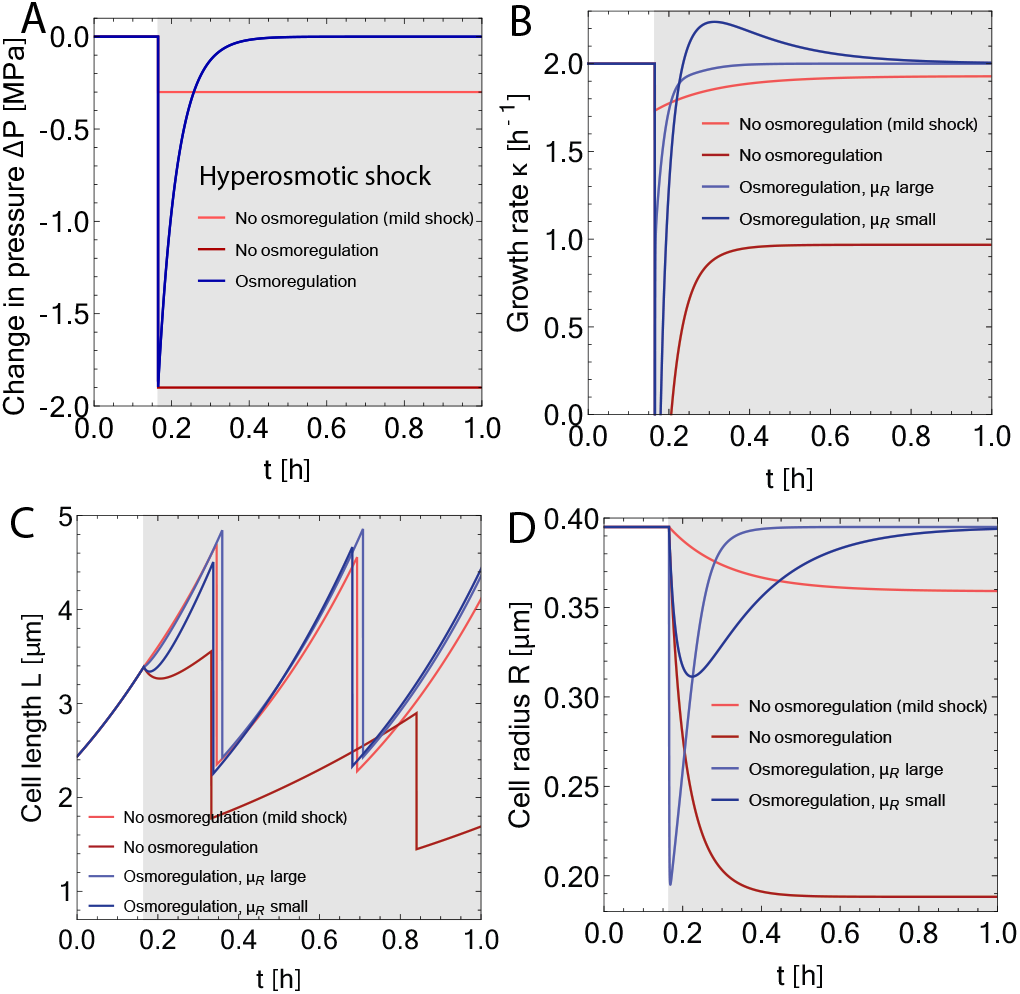
Adaptation to hyperosmotic shocks. (A) Change in turgor pressure (Δ*P* = *P*_0_ *− P*_*f*_) vs time plots under a hyperosmotic shock. The white region indicates the pre-shock period and the grey region indicates the post-shock environment. Osmoregulation is modeled here as an exponential relaxation of turgor pressure back to the preshock pressure *P*_0_ = 0.3 MPa with a decay constant 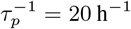. (B) Growth rate vs time plots of corresponding to the pressure perturbations in panel A. (C) Representative cell length trajectories corresponding to the pressure perturbations in panel A. (D) Cell radius vs time plots corresponding to the pressure perturbations in panel A.

To explain turgor pressure and cell shape recovery post hyperosmotic shocks, we examined a model of osmoregulation where turgor pressure relaxes back to its steady-state value *P*_0_ according to the equation:

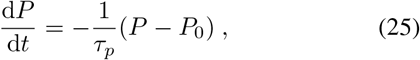

where *τ*_*p*_ is the time constant for relaxation. This model draws inspiration from the fact that bacterial cells possess a diverse set of osmoregulatory proteins that regulate turgor pressure homeostasis [60]. If cells undergo osmoregulation during the adaptation to osmotic shocks [61], both the radius and growth rate recover to their respective pre-shock values as *P* returns to *P*_0_. Additionally, if the timescale of radius changes (determined by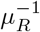) is sufficiently larger than the timescale of osmoregulation, the period during which pressure has recovered but radius has not corresponded to an overshoot in growth rate (Fig. 6B), consistent with experimental observations [23]. Our model thus predicts that the over-correction in growth rate following hyperosmotic shocks is a result of the mismatched timescales of pressure and morphological changes.

### B. Hypoosmotic shocks

In the case of hypoosmotic shocks, while small step function increases in pressure cause increases in growth rate (Fig. 7A-D), there is no growth rate recovery as is seen for hyperosmotic shocks. On the other hand, large pressure perturbations yield unexpected effects: cells initially shrink rather than swell and can reach new steady-state growth rates that are negative. This arises from the dominance of the *P* ^2^ term in strain energy over the turgor pressure energy − *PV* for large values of the final turgor pressure *P*_*f*_ . In other words, the cost of increased cell size can outweigh the minimization of energy from relieving pressure in the model. Even when pressure naturally relaxes due to volume changes and osmoregulation (Fig. 7A), the observed behavior resembles that of a hyperosmotic shock, with an initial sharp decline in growth rate and cell size followed by an overshoot in growth rate to the new steady-state (Fig. 7B-D). The value of *P*_*f*_ chosen for these simulations is large enough that the growth rate stabilizes to a negative value, indicating an unstable state within our model where the cell would either die or the optimal growth flux assumption is no longer valid.

**FIG. 7.**
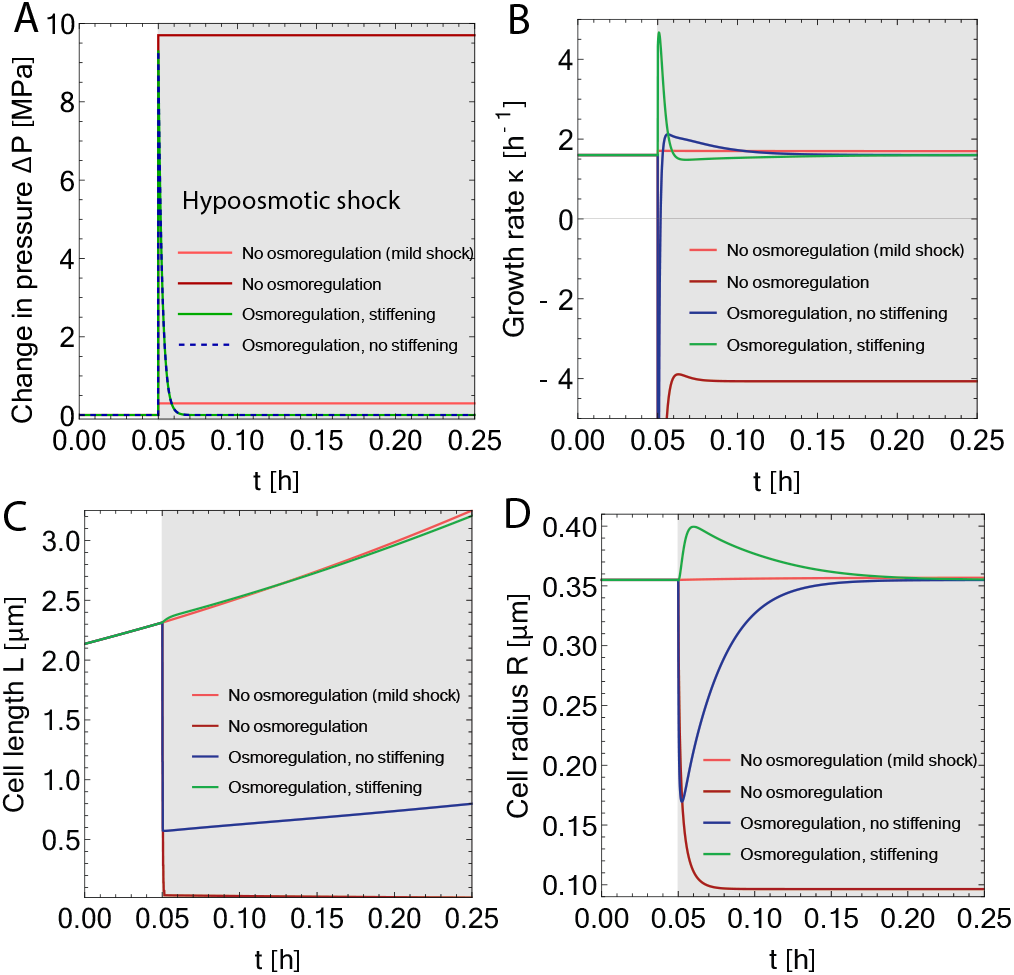
Adaptation to hypoosmotic shocks. (A) Change in turgor pressure (Δ*P*) vs time plots for a bacterial cell under a hypoosmotic shock. The white region indicates the pre-shock period and the grey region indicates the post-shock environment. Osmoregulation is modeled here as exponential decay of the post-shock turgor pressure back to the pre-shock pressure *P*_0_ = 0.3 MPa with a decay constant 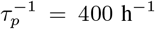. Temporary stifening of the cell envelope is also modeled as exponential decay from the perturbed value *λ/P* ^2^ = 0.23 MPa^−1^h^−1^ back to the pre-shock value with a decay constant 50 h^−1^. (B) Growth rate vs time plots corresponding to the pressure perturbations in panel A. (C) Representative cell length trajectories corresponding to the pressure perturbations in panel A. (D) Cell radius vs time plots corresponding to the pressure perturbations in panel A.

To more realistically model cellular response to hypoosmotic shocks within our theoretical framework, we need to consider potential changes in the mechanical properties of the cell wall. Recent work has both measured and modeled the softening of the cell wall in response to hyperosmotic shocks [59]. This prompts us to explore alterations in cellular strain energy under hypoosmotic shocks, which is governed by the parameter *λ* = *P* ^2^*ρ*_*g*_*/*4*k*_*s*_*ρ*_*p*_ (see Eq. 5). In contrast to hyperosmotic shocks, if the cell envelope stifens under hypoosmotic shock (increased *k*_*s*_) as it is stretched beyond the usual regime, *λ* would be expected to temporarily decrease. This decrease in *λ* could also result from a reduction in the glycan strand density (*ρ*_*g*_) as the cell temporarily swells at a greater rate than the synthesis of glycan strands.

We thus dynamically increased cell-wall stifness *k*_*s*_ during a hypoosmotic shock, using the same exponential relaxation form as in Eq. (25), with an independent decay constant. As shown in Fig. 7A-D, incorporating such a temporary increase in stifness *k*_*s*_ simultaneously with the increase in *P* results in the initial swelling of the cell as expected. This is followed by a relaxation of growth rate and cell sizes to a similar value as the pre-shock rate, with the potential for an undershoot in growth rate depending on *μ*_*R*_. The value of *P*_*f*_ chosen for this figure generates growth-rate trajectories that resemble the scale of the experimental data [23], indicating that the pressures required for the osmoregulation model to match data would not be stable growth conditions for non-osmoregulating cells. We thus predict that a temporary stiffening of the cell wall is essential to capture the experimentally observed growth rate dynamics [23] in response to a hypoosmotic shock. This prediction could be tested in future experiments.

## VI. DISCUSSION

All living organisms rely on energy assimilation and the efficient allocation of acquired energy to perform various physiological tasks. While theories connecting energy, metabolism, and growth have been developed for birds and mammals [1– 5], understanding energy allocation strategies in microorganisms has remained an open problem. In this paper, we address this gap by developing a theoretical framework for energy allocation in growing bacterial cells based on optimizing the rate of energy used for growth. We partition the use of imported energy into cellular metabolism and energy storage, with the former further divided into energy for growth, division, shape maintenance, and dissipation. Optimizing the energy flux to maximize growth results in equations of motion for cell size and protein production, predicting an increase in cell size with growth rate. While most of the model parameters can be directly estimated from published data on *E. coli*, a few are calibrated from experimental conditions or are determined by fitting the model to perturbation experiments.

Analyzing the energy dynamics throughout a cell cycle, we observe that the rate at which each energy sector accumulates energy is directly proportional to cell size. Across various growth environments, the fraction of energy utilized for mechanical tasks exhibits a minimum at a preferred cell size, while growth energy and dissipation increase with nutrient quality, trading off with stored energy and division. Notably, the fraction of energy allocated to growth increases with growth rate, consistent with the observation that the ribosome mass fraction also increases with growth rate [10].

An important implication of our theory is that it provides a rationale for the nonlinear increase in metabolic rate with cell size in bacteria [21]. Without incorporating additional regulatory mechanisms, our model predicts a superlinear scaling of metabolic rate with cell size, with a tunable scaling exponent determined by changes in stored energy density *g*. This superlinear metabolic scaling arises from the increase in cell size with growth rate. If cell size were independent of growth rate, a linear scaling of metabolic rate with cell size would be expected. Conversely, a decrease in growth rate with size would result in sublinear metabolic scaling, as observed in higher animals.

Given the dynamic formulation of our model, it is well suited to capture cellular behaviors out of steady state, such as during nutrient shifts. In nutrient upshifts, we find that nutrient import and cell size both steadily increase together during the shift, whereas in nutrient downshifts we can capture the observed undershoot in growth rate due to nutrient importation decreasing on a faster timescale than cell size.

The predicted behaviors of our model for nutrient shifts align with proteome partitioning models, where changes in growth rate are driven by shifts in proteome allocation, translation rate, and amino acid precursor levels [9, 15, 18]. A valuable comparison can be drawn between the energy allocation framework and proteome allocation models. In the latter, dynamics of proteome reallocation aim to maximize growth rate in the new environment, akin to our model where energy reallocation is dictated by the maximization of energy flux for growth, directly proportional to the growth rate. Although we do not explicitly model the allocation of ribosomal proteins that increases with nutrient quality, the increase in energy allocation to growth could be interpreted as originating from an increased mass fraction of ribosomal protein sector. Furthermore, the decrease in energy allocation to the division sector with growth rate is consistent with a reduced allocation to the division protein sector in proteome-centric models [18, 42]. The increase in nutrient import with growth rate corresponds to an elevated flux of amino acid precursors, resulting in a higher amino acid mass fraction during faster growth. From a proteome theory perspective, the observed growth rate undershoots in nutrient downshifts may be explained by a temporary undershoot in the amino acid mass fraction, as precursors are depleted faster than they are imported. This interpretation aligns with our model, where nutrient import decreases rapidly while cell size and associated energy costs lag behind.

One of the advantages of using the energy-based model as opposed to a proteome partitioning theory is that we can investigate the cellular response to mechanical perturbations without adding any new components to the model. By perturbing the turgor pressure, we can effectively capture the cellular response to hyperosmotic shocks with partial growth rate recovery and capture complete recovery if osmoregulation is taken into account. We find that osmoregulation is necessary to model hypoosmotic shocks, as well as changes to the mechanical properties of the cell wall. Recent work has elucidated that Gram-positive cells employ a feedback loop to sustain growth homeostasis following osmotic shocks [58]. As pressure changes cause expansion or contraction of the cell wall, altering membrane tension, this tension inversely influences precursor flux, thereby regulating growth rate to maintain homeostasis between wall and membrane synthesis. Although our model inherently includes feedback between mechanical stress, growth, and cell size, we currently lack feedback between nutrient import and mechanical stress in the cell envelope. To accurately model osmotic shock behavior in Gram-positive cells, it might be essential to integrate such a feedback system into our model.

Beyond addressing the cellular response to nutrient or osmotic stresses, our model holds the potential to investigate various environmental perturbations, including temperature changes [62] and stresses induced by antibiotics affecting cell growth and cell envelope properties [63–66]. The theory is not constrained by the shape of the cells [20] and the equations of motion can be easily developed for spherical or curved bacteria as well as cells with non-uniform shapes. This adaptability enables the application of the model to a diverse array of cell morphologies, such as flattened bacteria experiencing mechanical stress [67]. Moreover, recent experimental findings [68] revealing non-monotonic changes in ATP concentration between cell birth and division suggest a promising avenue for refining our understanding of energy utilization throughout the cell cycle by establishing connections between metabolic energy and ATP production.

## ACKNOWLEDGEMENTS

SB acknowledges support from the Royal Society (RGF/EA/181044), the National Institutes of Health (NIH R35 GM143042), and the Shurl and Kay Curci Foundation.

## Notes

### Competing Interest Statement

The authors have declared no competing interest.

